# *Bifidobacterium longum* attenuates ovariectomy-induced bone loss via modulating the Immunoporotic “Breg-Treg-Th17” cell axis

**DOI:** 10.1101/2022.01.13.476144

**Authors:** Leena Sapra, Niti Shokeen, Konica Gupta, Chaman Saini, Asha Bhardwaj, Mary Mathew, Pradyumna K. Mishra, Naibedya Chattopadhyay, Bhupendra Verma, Rupesh K. Srivastava

**Author notes:** Correspondence &.

## Abstract

Discoveries in the last few years have emphasized the existence of an enormous breadth of communication between osteo-immune system. These discoveries fuel novel approaches for the treatment of several bone-pathologies including osteoporosis, an inflammatory bone anomaly affecting more than 500 million people globally. *Bifidobacterium longum* (BL) is preferred probiotic of choice due to its varied immunomodulatory potential in alleviating various inflammatory diseases. Here, we evaluate the effect of BL in ovariectomy (ovx)-induced post-menopausal osteoporotic mice model. Our in vitro findings reveal that BL suppresses the differentiation and functional activity of RANKL-induced osteoclastogenesis in both mouse bone marrow cells and human PBMCs. Our in vivo data clearly establish that BL exhibits osteoprotective potential via modulating the “immunoporotic” Breg-Treg-Th17 cell-axis. Furthermore, µCT and bone mechanical strength data support that BL supplementation significantly enhanced bone mass and strength, and improved microarchitecture in ovx mice. Remarkably, alteration in frequencies of CD19^+^CD1d^hi^CD5^+^ Bregs, CD4^+^Foxp3^+^IL-10^+^ Tregs, and CD4^+^Rorγt^+^IL-17^+^ Th17 immune cells in distinct lymphoid organs along with serum-cytokine data (enhanced anti-osteoclastogenic cytokines IFN-γ and IL-10 and reduced osteoclastogenic-cytokines IL-6, IL-17, and TNF-α) strongly support the immunomodulatory potential of BL. Altogether our findings establish a novel osteo-protective and immunoporotic potential of BL in augmenting bone health under osteoporotic conditions.

## Introduction

Osteoporosis is defined as a systemic bone loss disease exemplified by deterioration of bone tissues and low bone mineral density (BMD), subsequently leading to fragility-related fractures. Osteoporosis is considered as a serious public health issue accounting for a huge socioeconomic burden. Post-menopausal osteoporosis mainly stems from the cessation of ovarian function and decline in estrogen that stimulates bone resorption and rapid bone loss ^1^. Although, currently available therapies to treat osteoporosis are effective, side effects (real and perceived) are impediment to treatment adherence by the patients. Various dietary supplements hold promise in conserving bone in postmenopausal condition.

“Immunoporosis” is an emerging area of research that studies the involvement of immune system in osteoporosis^2^. In this regard, the roles of regulatory T cells (Tregs) and inflammatory T cells (Th17) in bone homeostasis have been amply elucidated. Recently, our group along with others have reported that the imbalance between Treg-Th17 cells are key to the pathophysiology of bone-related diseases including osteoporosis ^2,3^. Th17 cells are established osteoclastogenic subset of T cells owing to the production of osteoclastogenic cytokines including IL-6, TNF-α, IL-17, and RANKL ^4,5^. On the contrary, Tregs are anti-osteoclastogenic T cell subsets that produce anti-osteoclastogenic cytokines including IL-10 and TGF-β. CD4^+^CD25^+^Foxp3^+^ Tregs suppress osteoclastogenesis and bone resorptive functions of osteoclasts via TGF-β1 and IL-10 in human peripheral blood mononuclear cells (PBMCs) ^6^. Recently, we demonstrated that Bregs suppressed osteoclastogenesis in vitro and protected ovx mice from bone loss thus underscoring the importance of this subset of cells in mitigating osteoporosis^7^.

Bregs also modulate the differentiation of Tregs and Th17 cells in autoimmune diseases including type 1 diabetes (T1D), systemic lupus erythematosus (SLE), and rheumatoid arthritis (RA)^8^. A recent study showed that the adoptive transfer of Bregs into osteoporotic mice successfully reduced the deterioration in alveolar bone by lowering the percentage of IL-17-producing Th17 cells ^9^. Thus, pharmacological modulation of the Bregs-Tregs-Th17 cell axis could represent a novel bone anabolic therapy for osteoporosis.

In recent years, the “Gut-Immune-Bone” axis has gained much attention from researchers worldwide. In humans, approximately 100 trillion microbes reside with the major fraction residing in the gastrointestinal tract. According to the WHO guidelines, probiotics are viable micro-organisms that impart beneficial health effects when administered in adequate amounts. Various evidences have demonstrated that manipulation of the gut microbiota via administration of probiotics enhance bone heath^10–12^. One of the widely studied multifunctional probiotics strains of *Bifidobacterium* species is *Bifidobacterium longum* (BL). It is a gram-positive, anaerobic bacterium and among the first microbes to colonize the human gastrointestinal (GI) tract and modulates the entire microbial diversity. BL is found to be effective in alleviating gastrointestinal and infectious diseases by stabilizing the gut microbiota and intestinal environment ^13^. A study reported that BL administration enhanced the BMD of ovx rats by upregulating the expression of bone morphogenetic protein-2 (BMP-2) gene ^14^. *Bifidobacterium* also alters the gut microbiota thereby modulating the metabolism of Tregs ^15^. However, there is no study that defined the immunomodulatory role of BL and its impact on augmenting bone health in osteoporosis has not been investigated till date. Thus, in the present study, we investigated the immunoporotic potential of BL in enhancing bone health in the post-menopausal osteoporotic (ovx) mice model.

To our knowledge, this is the first study that reveals the osteo-protective role of BL in ovx mice model via its ability to regulate the immunomodulatory “Bregs-Tregs-Th17” cell axis. Our both in vitro and in vivo data strongly suggest that BL exhibits the potential of suppressing osteoclastogenesis via enhancing the osteoprotective potential of Bregs. Of note, the immunomodulatory potential of BL is further strengthened from our serum cytokine data where we observe enhanced levels of anti-osteoclastogenic cytokines i.e., IL-10 (signature cytokine of Breg and Tregs) and IFN-γ together with reduced levels of osteoclastogenic cytokines i.e., TNF-α, IL-6, and IL-17 (signature cytokine of Th17 cells). Altogether, the present study underlines the osteoprotective role of BL via modulating the Immunoporotic “Breg-Treg-Th17 cell axis” thereby opening novel avenues both for the treatment and management of inflammatory bone loss observed in postmenopausal osteoporosis.

## Results

### BL suppress differentiation and function of osteoclasts

To elucidate the involvement of BL in modulating bone health, we initially examined the potential of BL on receptor activator of nuclear kappa B ligand (RANKL) induced osteoclasts differentiation and their functional activity. A schematic diagram presents the experimental protocol (Fig 1A, and for details refer to the Methods section). Bone marrow cells (BMCs) were stimulated with osteoclastogenic media supplemented with M-CSF (30 ng/ml) and RANKL (100 ng/ml) in the presence or absence of BL-CM at different ratios (1:100, 1:10, and 1:5). After 4 days, cells were harvested and processed for both TRAP staining and F-actin ring polymerization assay. There was a significant reduction in the differentiation of osteoclasts in a dose-dependent manner as evident by a significant reduction in multinucleated (> 3 nuclei) TRAP-positive cells in BL treated group in comparison to control group (Fig. 1B-D). Furthermore, the area of multinucleated osteoclasts in treatment groups was significantly less than the control group (Fig. 1E). Also the number and the area of F-actin rings upon BL treatment were significantly decreased upon BL treatment, which indicated that BL not only inhibited osteoclastogenesis but also the functional activity of osteoclasts (Fig. 1F-I).

**Figure 1:**
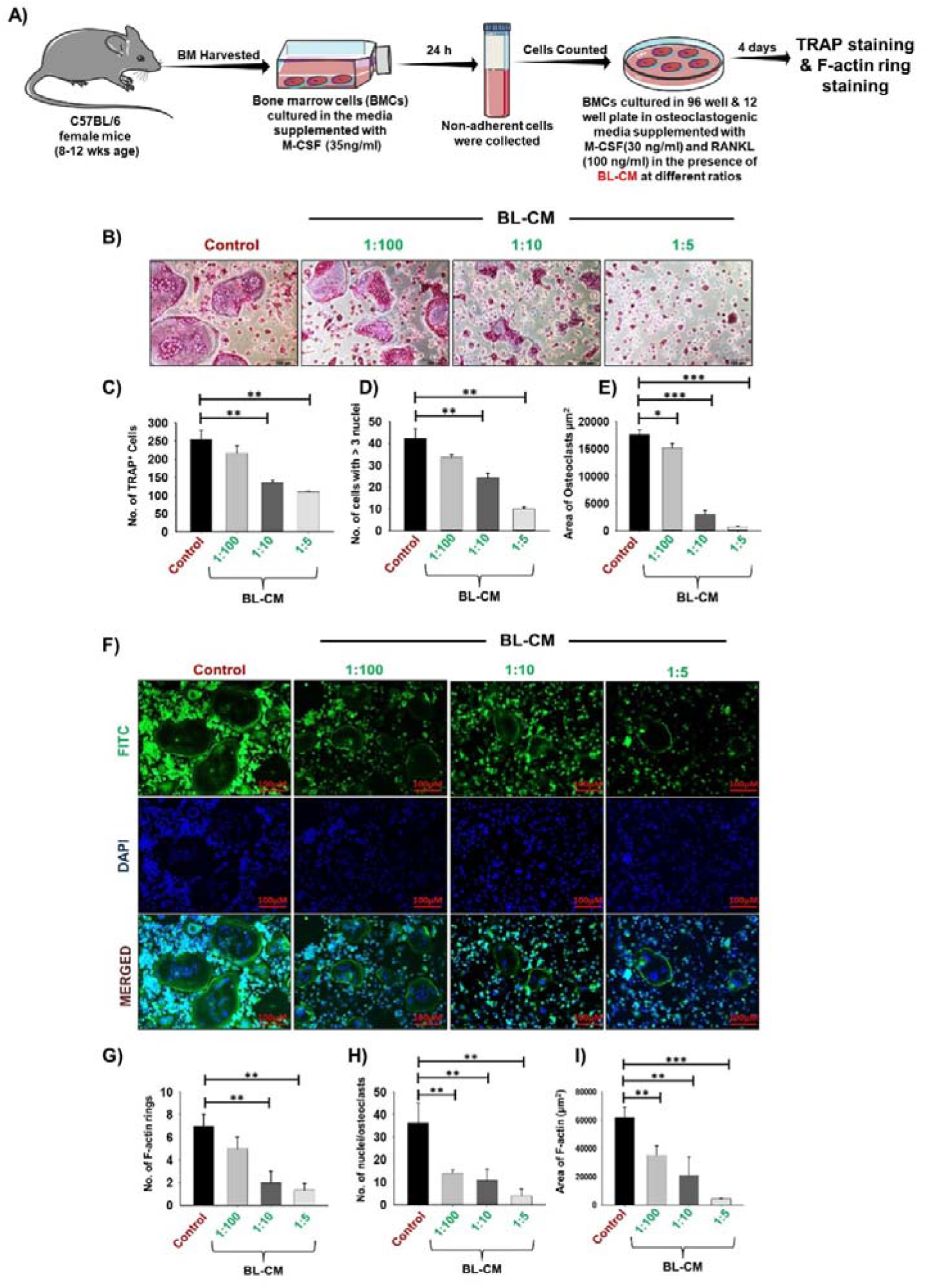
BL suppress osteoclastogenesis and F-actin polymerization in a dose dependent manner: A) Osteoclast differentiation was induced in Bone Marrow cells (BMCs) with M-CSF (30 ng/ml) and RANKL (100 ng/ml) with or without *Bifidobacterium longum*—conditioned media (BL-CM) at different ratios of 1:100, 1:10 and 1:5 for 4 days. Giant multinucleated cells were stained with TRAP and cells with ≥ 3 nuclei were considered as mature osteoclasts. B) Photomicrographs at 20X magnification were taken. C) Number of TRAP positive cells; D) Number of TRAP positive cells with more than 3 nuclei. E) Area of osteoclasts. F) F-actin and nuclei were stained with FITC-conjugated phalloidin and DAPI respectively. Images were captured in fluorescence microscope (Imager.Z2 Zeiss microscope) at 10× magnification. G) Number of F-actin rings. H) Number of nuclei per osteoclasts. I) Area of F-actin ring. The above images are indicative of one independent experiment and similar results were obtained in atleast three different independent experiments. Statistical significance was considered as p ≤ 0.05 (*p ≤ 0.05, **p ≤ 0.01, ***p ≤ 0.001) with respect to indicated groups.

### BL possess immunomodulatory potential

Immune cells play a vital role in protection against pathogenic organisms and in maintaining the homeostatic bone remodeling processes ^2^. Besides, it has been observed that the adaptive immune system is overwhelmingly shaped by the gut microbial species ^16^. Among various subsets of T lymphocytes, Treg and Th17 cells are important modulators of the bone remodeling process ^2,3^. Thus, we next investigated the potential of BL in regulating Th17 and Treg immune cell balance for which, naïve T cells were differentiated in Th17 and Treg polarizing conditions in the absence or presence of BL-CM at distinct ratios (1:100, 1:10, and 1:5) (Fig. 2A). At the end of incubation, flow cytometry was performed for analyzing the percentages of CD4^+^Foxp3^+^IL-10^+^ Tregs and CD4^+^Rorγt^+^IL-17^+^ Th17 cells. Notably, we found that treatment with BL-CM significantly reduced the percentages of Th17 cells (4-fold) and IL-17 cytokine production (2-fold) (Fig. 2B-F). We also observed that BL-CM treatment significantly enhanced the percentages of both Type1 regulatory T (Tr1) cells (5-fold) and Tregs (2-fold) as well as their efficiency to express IL-10 cytokine (4-fold) (Fig. 2G-M). These results suggest the potential of BL in modulating Th17-Treg cell balance. Of note, several studies revealed that naïve T cells differentiation into Tregs and Th17 cells is further regulated by Bregs under physiological conditions ^17^. Hence, we estimated whether BL modulated the differentiation of B cells into Bregs. For this purpose, negatively selected splenic B cells were stimulated under Breg polarizing conditions in the presence of BL-CM for 24 h (Fig. 3A). We observed that BL significantly enhanced the percentage of CD19^+^CD1d^hi^CD5^+^ Bregs (p < 0.05) along with a significant increase in IL-10 production (Fig. 3B-E). Taken together, these results indicate the immunomodulatory roles of BL.

**Figure 2:**
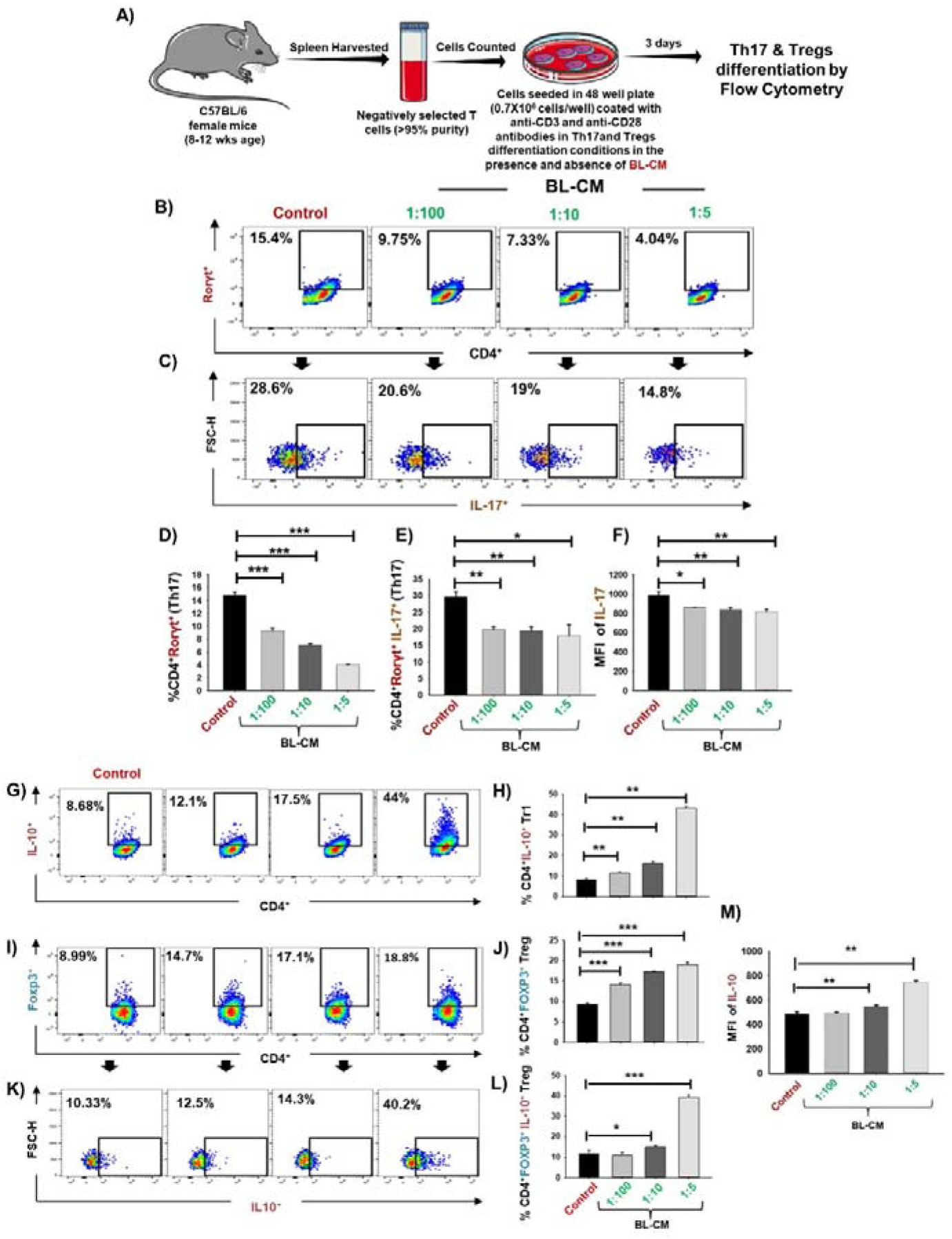
BL inhibits Th17 cell differentiation and enhances Tr1 and Treg differentiation in vitro: A) For Th17 and Treg cell differentiation, splenic naïve CD4^+^CD25^-^ T cells stimulated with anti-CD3 and anti-CD28 mAbs were incubated with differentiation cytokines in the presence and absence of BL-CM at different ratios. After 3 days, cells were analyzed for Th17, Tr1 and Treg differentiation. B) Dot plots depicting the percentages of CD4^+^Rorγt ^+^ Th17 cells. C) Dot plots depicting the percentages of CD4^+^ Rorγt ^+^IL-17^+^ Th17 cells. D) Graphical representation of percentage of CD4^+^Rorγt^+^Th17 cells. E) Percentage of CD4^+^Rorγt^+^IL-17^+^ Th17 cells. F) Mean fluorescence intensity (MFI) of IL-17. G) Dot plots and bar graphs depicting the percentages of CD4^+^ IL-10^+^ Tr1 cells. H) Graphical representation of percentage of CD4^+^IL-10^+^ Tr1 cells. I) Dot plots depicting the percentages of CD4^+^ FOXP3^+^ Tregs. J) Bar graphs depicting the percentages of CD4^+^FOXP3^+^ Tregs. K) Dot plots depicting the percentages of CD4^+^FOXP3^+^IL-10^+^ Tregs. L) Bar graphs depicting the percentages of CD4^+^FOXP3^+^IL-10^+^ Tregs. M) Mean fluorescence intensity of IL-10 cytokine. Data is represented as Mean ±SEM. Similar results were obtained in atleast different independent experiments. Statistical significance was considered as p ≤ 0.05 (*p ≤ 0.05, **p ≤ 0.01, ***p ≤ 0.001) with respect to indicated groups.

**Figure 3:**
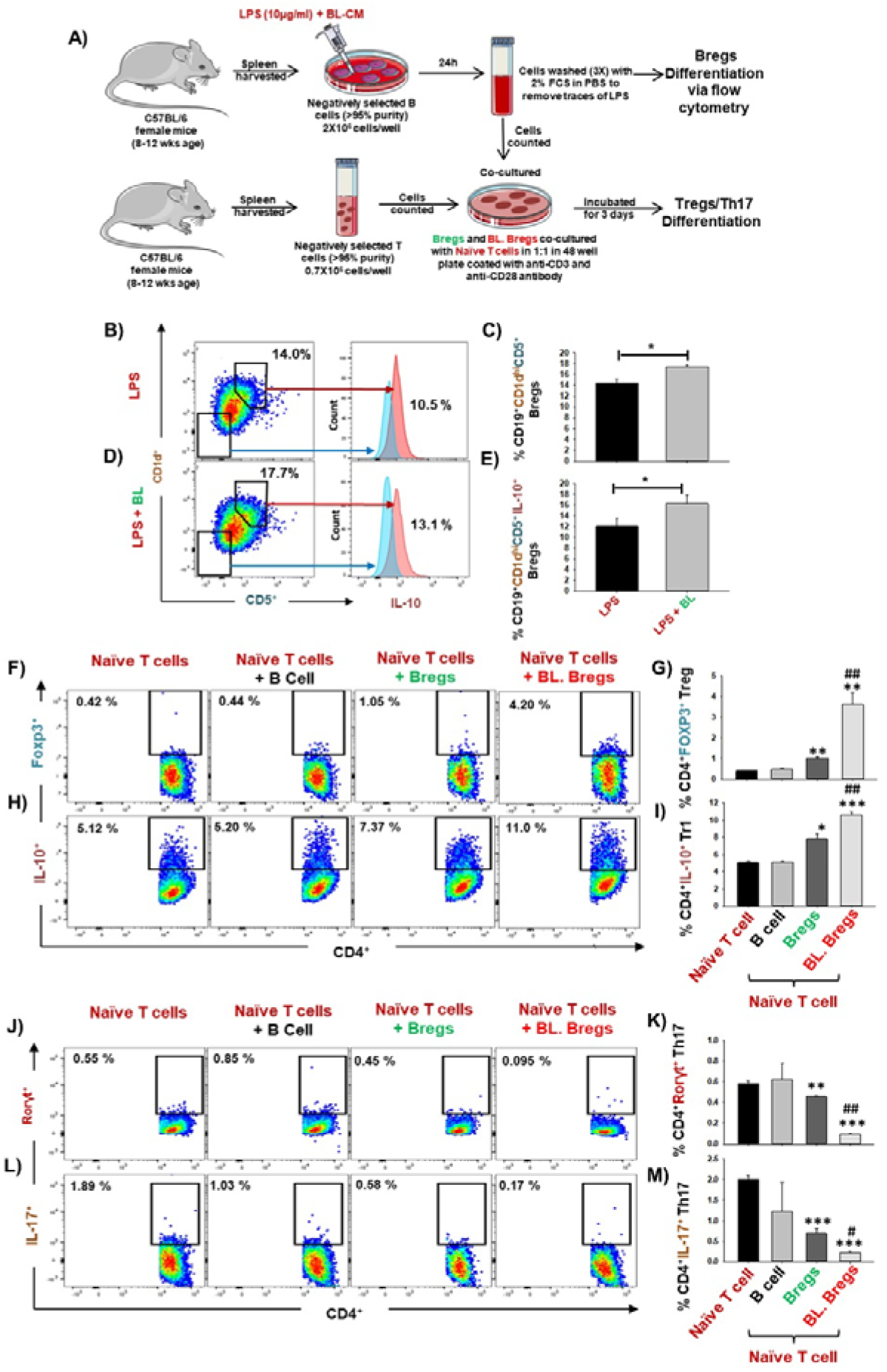
BL enhances immunosuppressive potential of Bregs: A) For Bregs stimulation, splenic CD19^+^ B cells were negatively selected and stimulated with LPS (10ug/ml) in the presence and absence of BL-CM at 1:5 dilution. After 24 hours, cells were analyzed for CD1d, CD5 and IL-10 expression by FACS. B) Dot plots depicting the percentages of CD19^+^CD1d^high^CD5^+^ Bregs. C) Bar graphs representing the percentages of CD19^+^CD1d^high^CD5^+^ Bregs. D) Dot plots depicting the percentages of CD19^+^CD1d^high^CD5^+^IL-10^+^Bregs. E) Bar graphs representing the percentages of CD19^+^CD1d^high^CD5^+^IL-10^+^Bregs. F) Naïve T cells and LPS stimulated and LPS + BL-CM were co-cultured in anti-CD3 and CD28 coated plate for 3 days. B cells were induced with LPS (10 ug/ml) and BL-CM (1:5) for 24h prior to co-cultures. Dot plots depicting the percentages of CD4^+^FOXP3^+^ Tregs. G) Bar graphs representing the percentages of CD4^+^FOXP3^+^ Tregs. H) Dot plots depicting the percentages of CD4^+^IL-10^+^ Tr1 cells. I) Bar graphs representing the percentages of CD4^+^IL-10^+^ Tr1 cells. J) Dot plots depicting the percentages of CD4^+^Rorγt^+^Th17 cells. K) Bar graphs representing the percentages of CD4^+^Rorγt^+^Th17 cells. L) Dot plots depicting the percentages of CD4^+^IL-17^+^ Th17 cells. M) Bar graphs representing the percentages of CD4^+^IL-17^+^ Th17 cells. (* denotes comparison of indicated group with respect to naïve T cells and # denotes comparison of indicated group with respect to Bregs group). Data is represented as Mean ±SEM. Similar results were obtained in atleast three different independent experiments. Statistical significance was considered as p ≤ 0.05 (*p ≤ 0.05, **p ≤ 0.01, ***p ≤ 0.001) with respect to indicated groups.

### BL-stimulated Bregs regulate Treg-Th17 cell balance

Moving ahead in our study, we were next keen in assessing the immunomodulatory potential of BL-induced Bregs in regulating the differentiation of naïve T cells either into Tregs or Th17 cells under non-polarizing conditions. To this aim, we co-cultured BL stimulated Bregs with negatively selected naïve T cells at a 1:1 ratio for 3 days (Fig. 3A) in anti-CD3 and anti-CD28 coated plates under non-polarizing conditions. Subsequently, cells were harvested and analyzed for the percentages of CD4^+^Foxp3^+^IL-10^+^ Tregs and CD4^+^Rorγt^+^IL-17^+^ Th17 cells by flow cytometry. We observed that BL-stimulated Bregs significantly increased the percentage of both CD4^+^Foxp3^+^IL-10^+^ Tregs (p < 0.01) and CD4^+^IL-10^+^ Tr1 cells (p < 0.001) in comparison to control groups (Fig. 3F-I). Besides, BL-stimulated Bregs significantly reduced the percentage of CD4^+^Rorγt^+^IL-17^+^ Th17 cells (p < 0.001) (Fig. 3J-M). Taken together, these data suggests that the enhancement in the percentage of Bregs in response to BL stimulation is pivotal for the efficient induction of anti-osteoclastogenic Tregs along with concurrent inhibition of osteoclastogenic Th17 cells, thereby pointing towards role of “Breg-Treg-Th17” cell axis.

Recently, our group reported that Bregs exhibit the potential to inhibit osteoclasts differentiation ^7^. Thus, we studied whether BL modulated the anti-osteoclastogenic potential of Bregs. To this aim, we co-cultured BL-induced Bregs and BMCs at various ratios (10:1, 5:1, and 1:1) for 4 days (Fig. 4A). Our TRAP data showed that BL-induced Bregs inhibited the generation of osteoclasts more efficiently than the non-BL stimulated Bregs (Fig. 4B-D). These data indicate that BL could prevent bone loss by suppressing osteoclasts differentiation along with favourable modulation of the pivotal immunoporotic “Breg-Treg-Th17” cell axis.

**Figure 4:**
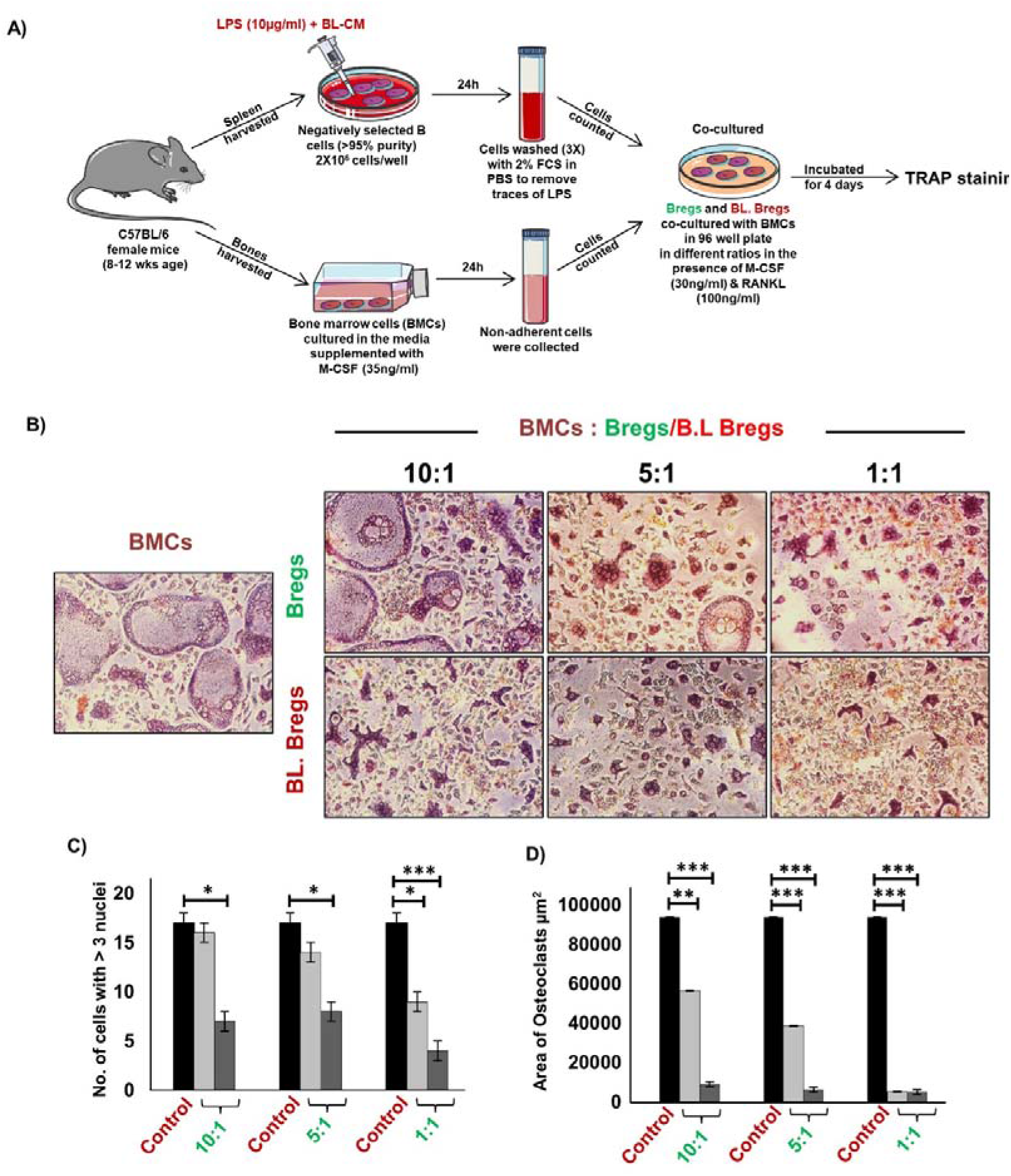
BL enhances anti-osteoclastogenic potential of Bregs: A) BMCs and LPS stimulated and LPS + BL-CM were co-cultured in cell culture plate in the presence of M-CSF (30 ng/ml) and RANKL (100 ng/ml) for 4 days. B cells were induced with LPS (10 ug/ml) and BL-CM (1:5) for 24h prior to co-cultures. B) LPS + BL-CM induced Bregs suppress osteoclastogenesis more efficiently in comparison to LPS induced Bregs. C) Number of TRAP positive cells with more than 3 nuclei. D) Area of osteoclasts. The above images are indicative of one independent experiment and similar results were obtained in atleast three different independent experiments. Statistical significance was considered as p ≤ 0.05 (*p ≤ 0.05, **p ≤ 0.01, ***p ≤ 0.001) with respect to indicated groups.

### BL ameliorates bone loss

We next assessed the effect of BL in mitigating bone loss caused by estrogen deficiency (ovx) condition using adult C57BL/6 mice that were distributed into three groups: sham surgery (ovary intact), bilateral ovx, and ovx group orally administered daily with BL (10^9^cfu) for 6 weeks. At the end of the treatments, mice were killed by euthanasia, and the bones were collected for further studies (Fig. 5A). There was no difference in the bodyweight between the groups throughout duration of the study (Fig. S1). Scanning electron microscopy (SEM) of the femoral cortical bone sections showed higher bone resorption pits or lacunae in the ovx group compared with the sham group whereas the BL group showed a significant mitigation of pits compared with the ovx group (Fig. 5B). To further examine these 2D SEM images quantitively in a more statistical manner we performed MATLAB analysis to obtain the association between bone loss and bone mass. MATLAB analysis of SEM images represents the magnitude of homogeneity where the higher correlation is denoted by red color (enhanced bone mass) whereas blue color (lower correlation) symbolizes enhanced bone loss. MATLAB analysis of 2D SEM images indicated that BL administered ovx group showed a higher correlation value and thus indicating enhanced bone mass (Fig. 5C). We next estimated the 3D topology of the bone surface in response to BL treatment by AFM which indicated reduced bone resorption and roughness of the bone surface in the BL administered ovx group (Fig. 5D). Moreover, MATLAB analysis of AFM images suggests an improved bone architecture and decreased osteoclastogenesis in BL group compared with the ovx group (Fig. 5E). These findings, strongly indicate that BL attenuates bone loss in ovx mice.

**Figure 5:**
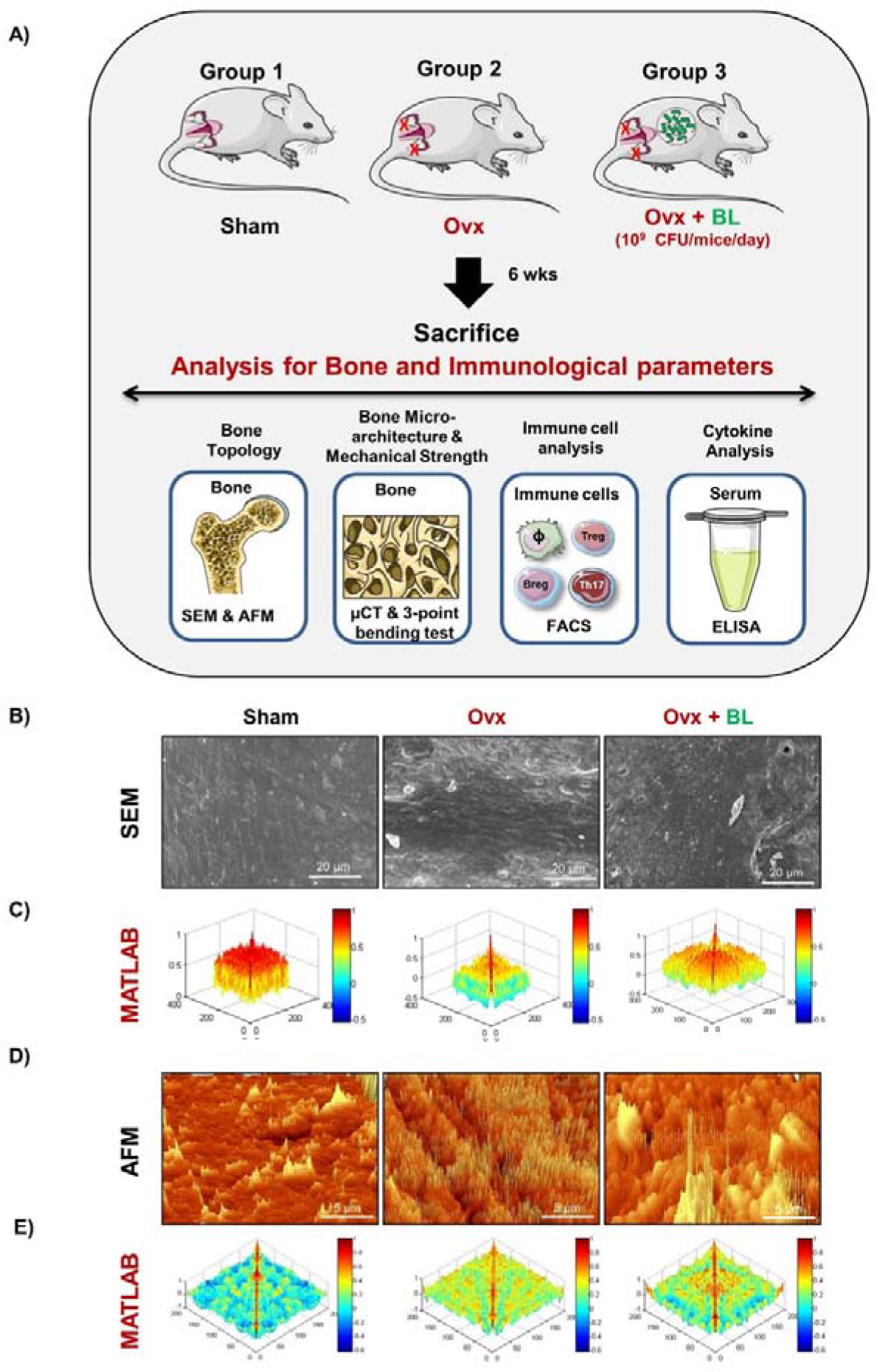
BL administration attenuates bone loss in Ovx mice: Experimental Layout followed for in vivo studies Mice were divided into 3 groups viz. Sham, Ovx and Ovx + BL group that received BL at 10^9^ CFU/day orally reconstituted in drinking water. At the end of 45 days, mice were sacrificed and analysed for various parameters. B) 2D SEM images C) 2D MATLAB analysis of SEM images D) 3D AFM images. E) 3D MATLAB analysis of AFM images. The representative images are indicative of one independent experiment and comparable results were obtained in two di erent independent experiments with n=6 mice/group/experiment.

### BL maintains bone microarchitecture in ovx mice

Osteoporosis and fracture risk is high in both spine and hip of postmenopausal women. These two anatomical regions are represented by lumbar vertebra (5^th^) and proximal femur metaphysis. Micro-computed tomography (µ-CT) assessment of lumbar vertebrae (LV)-5 showed loss of bone volume and deteriorated microarchitecture in the ovx mice compared with both sham and BL treated groups (Fig. 6A). BL supplementation significantly enhanced bone volume per tissue volume (BV/TV) (p < 0.01), trabecular thickness (Tb.Th) (p < 0.01); and decreased trabecular separation (Tb.Sp) (p < 0.05) compared with the ovx group (Fig. 6B).

**Figure 6:**
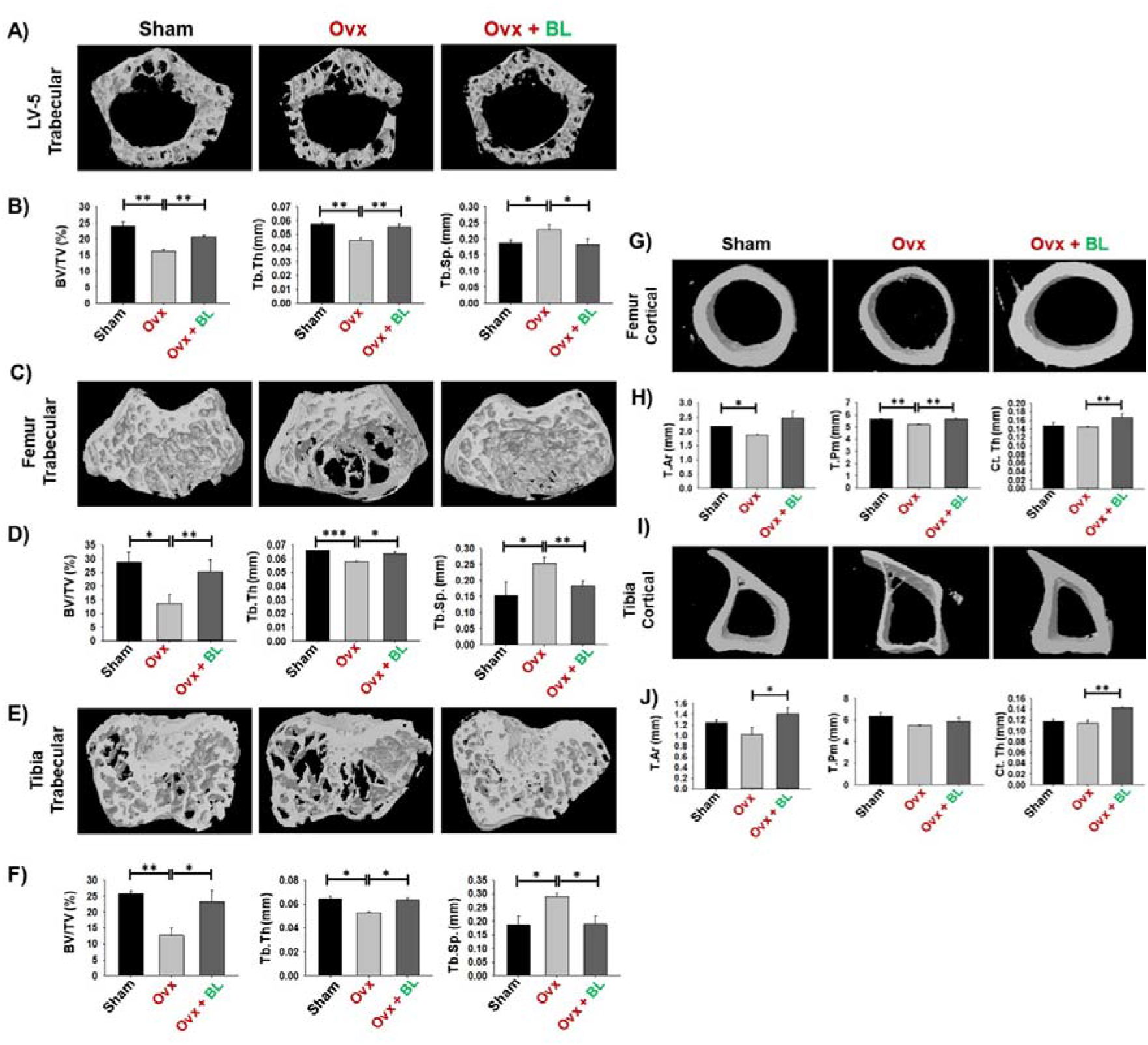
BL administration improves both trabecular and cortical bone microarchitecture. 3-D uCT reconstruction of LV-5 Trabecular, Femur Trabecular, Tibia Trabecular, Femur Cortical and Tibia Cortical of all groups. A) Bone micro-architecture of Femur trabecular. B) histomorphometric parameters of Femur trabecular. C) Bone micro-architecture of Tibia trabecular. D) Histomorphometric parameters of Tibia trabecular. E) Bone micro-architecture of LV-5 trabecular. F) Histomorphometric parameters of LV-5 trabecular. Histomorphometric parameters Bone volume/tissue volume ratio (BV/TV); Tb. Th., trabecular thickness; Tb. Sp., trabecular separation. G) Bone micro-architecture of Femur cortical. H) Histomorphometric parameters of Femur cortical. I) Bone micro-architecture of Tibia cortical. J) Histomorphometric parameters of Tibia cortical. Histomorphometric parameters of Tibia Cortical. Tt. Ar., total cross-sectional area; T. Pm., total cross-sectional perimeter; Ct. Th., cortical thickness. The results were evaluated by ANOVA with subsequent comparisons by Student t-test for paired or nonpaired data. Values are reported as mean ± SEM. The above graphical representations are indicative of one independent experiment and similar results were obtained in two different independent experiments with n=6. Statistical significance was considered as p≤0.05 (*p≤0.05, **p≤0.001, ***p≤0.0001) with respect to indicated mice groups.

We next investigated the effect of BL on appendicular bones including femur and tibia. 3D-micro-architecture of proximal metaphyses of femur and tibia showed that BV/TV and Tb.Th were decreased and Tb.Sp increased in ovx mice compared with sham, suggesting loss of trabecular bones; and BL treatment reversed all the ovx-induced changes (Fig. 6C-F). Cortical bones of femur and tibia showed thinning due to ovx as periosteal area (T.Ar), periosteal perimeter (T.Pm) and cortical thickness (Cs.Th) were decreased compared with the sham; and BL treatment reversed these changes (Fig. 6G-J). These data demonstrate that BL administration significantly improves 3D micro-architecture and histomorphometric parameters of both trabecular and cortical bones in osteoporotic mice,.

### BL enhance both bone mineral density and mechanical strength

Since BMD is a predictor of osteoporotic fracture, we measured it at all weight-bearing bones of axial and appendicular sites. μCT allows measurement of volumetric BMD and our data showed that it was significantly decreased at all sites measured in ovx mice compared with sham; and BL treatment reversed these changes (Fig. 7A-E).

**Figure 7:**
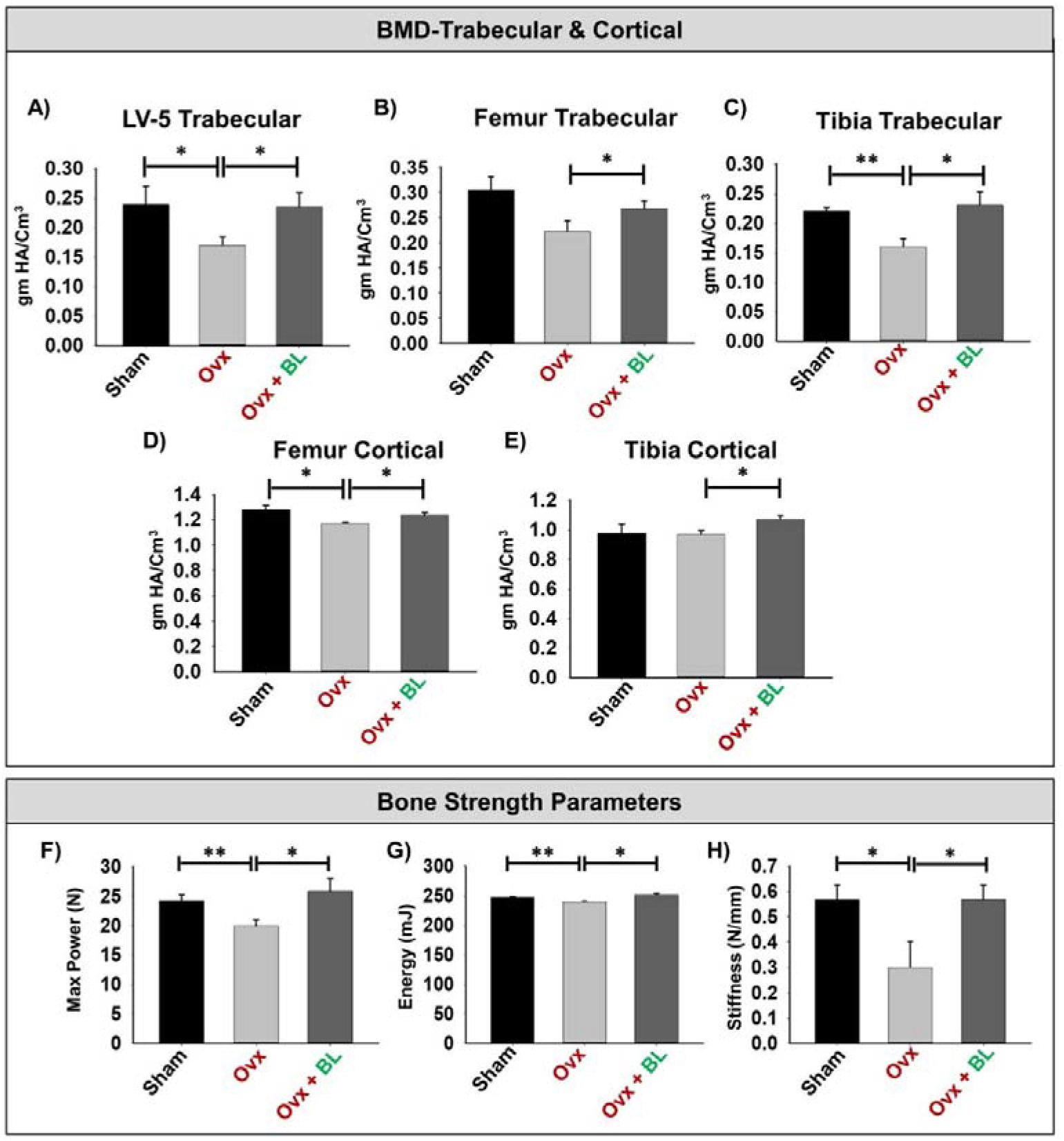
BL administration enhances Bone Mineral Density (BMD) and Mechanical strength of bones. A) Graphical representation of BMD of LV-5 trabecular region. B) Graphical representation of BMD of femur trabecular region. C) Graphical representation of BMD of tibia trabecular region. D) Graphical representation of BMD of femur cortical region. E) Graphical representation of BMD of tibia cortical region. F) Graphical representation of bone mechanical strength parameters Maximum power (N). G) Graphical representation of energy (mJ). H) Graphical representation of stiffness (N/mm). Data are reported as mean ± SEM. Similar results were obtained in two independent experiments with n=6. Statistical significance of each parameter was assessed by ANOVA followed by paired group comparison. *p < 0.05, **p < 0.01, ***p < 0.001, ****p < 0.0001 compared with indicated groups.

We next assessed the bone quality by measuring the bending strength of femurs. Maximum load, failure energy and stiffness were significantly decreased in the ovx mice compared with sham, and these parameters were maintained to the sham level in the BL group (Fig. 7F-H). These data suggest maintenance of bone mass and strength in the ovx mice treated with BL.

### BL promotes bone heath by modulating the immunoporotic “Breg-Treg-Th17” cell axis

Our in vitro data showed that BL has anti-osteoclastogenic potential achieved by the favourable modulation of the “Breg-Treg-Th17” cell axis. Next, we measured Breg, Treg, and Th17 cell population in lymphoid organs including bone marrow (BM), spleen, mesenteric lymph nodes (MLN) and Peyer’s patches (PP) in response to ovx and BL treatment. Ovx mice had higher percentages of CD4^+^Rorγt^+^ Th17 cells in BM (3-fold, p < 0.05), spleen (3.5-fold, p < 0.05), MLN (2-fold, p < 0.01) and PP (2-fold, p < 0.05) than the sham; and BL treatment reversed these changes (Fig. 8A-H). Conversely, the percentages of CD4^+^Foxp3^+^ Tregs in BM (1.5-fold, p < 0.05), spleen (2-fold, p < 0.05), MLN (p < 0.05) and PP (2-fold, p < 0.05) in the ovx group were lower than the sham; and BL treatment restored these parameters (Fig. 9A-H). As under physiological conditions the balance between Treg and Th17 cells is also regulated by Bregs, we next investigated whether the observed imbalance of Treg and Th17 cells was due to the alteration in the frequency of the Breg population. CD19^+^CD1d^hi^CD5^+^ Bregs population was significantly decreased in BM (p < 0.05) and spleen (p < 0.01) of ovx mice compared with sham; and BL administration to ovx increased this cell population to sham level (Fig. 9I-L). Besides, the circulating levels of pro-osteoclastogenic cytokines including IL-6, TNF-α and IL-17 were increased and anti-osteoclastogenic cytokines such as IFN-γ and IL-10 were decreased in ovx mice compared with sham. BL treatment of ovx mice completely reversed the cytokine profiles, which suggested its role in attenuating inflammatory bone loss in postmenopausal osteoporosis.

**Figure 8:**
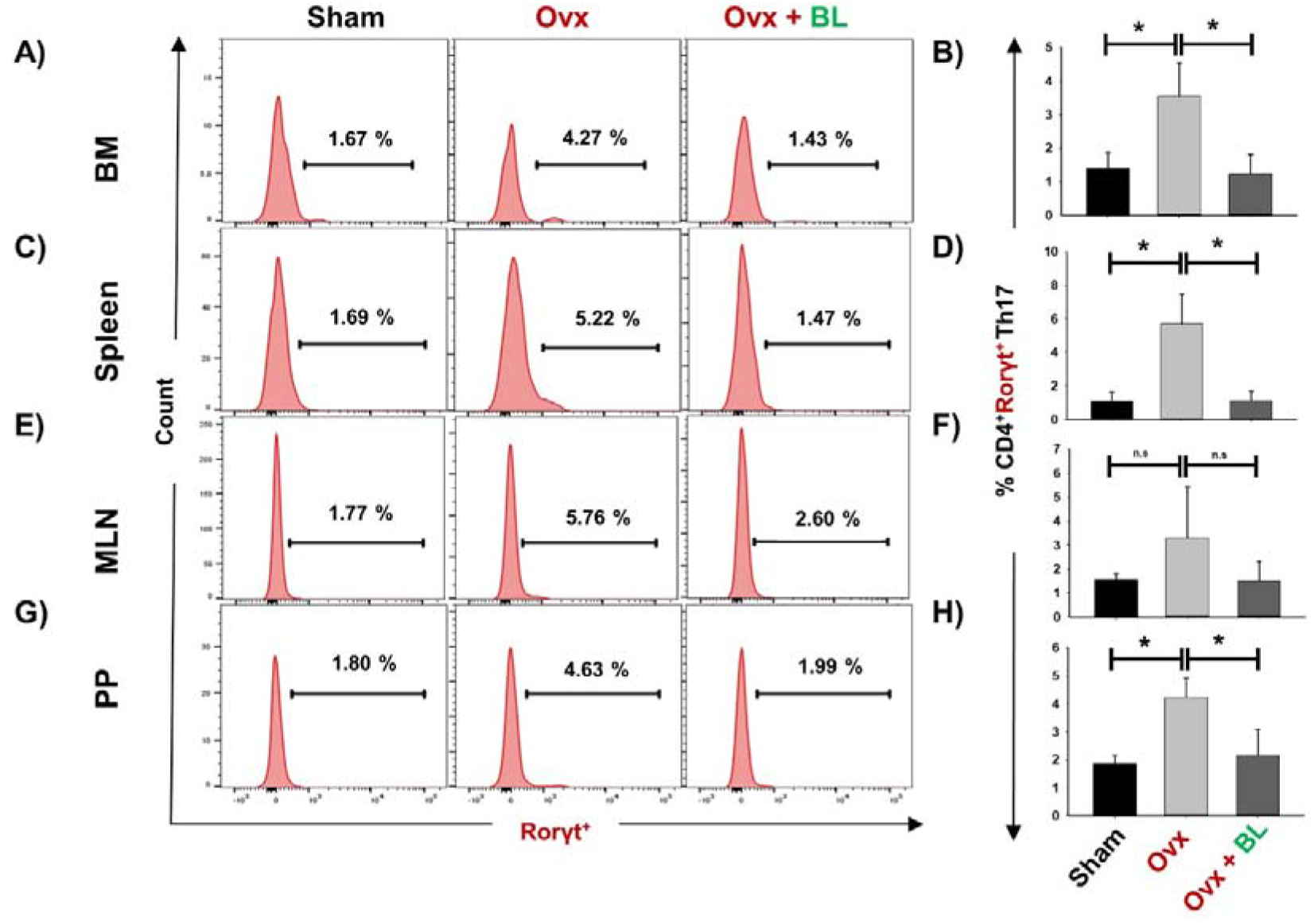
BL administration inhibits osteoclastogenic Th17 cells *in vivo*. Cells from various lymphoid organs were harvested and analysed for CD4^+^Rorγt^+^ Th17 cells. A) Histograms are representing percentages of CD4^+^Rorγt^+^ Th17 cells in BM. B) Bar graphs are representing percentages of CD4^+^Rorγt^+^ Th17 in BM. C) Histograms are representing percentages of CD4^+^Rorγt^+^ Th17 cells in spleen. D) Bar graphs are representing percentages of CD4^+^Rorγt^+^ Th17 in spleen. E) Histograms are representing percentages of CD4^+^Rorγt^+^ Th17 cells in MLN. F) Bar graphs are representing percentages of CD4^+^Rorγt^+^ Th17 in MLN. G) Histograms are representing percentages of CD4^+^Rorγt^+^ Th17 cells in PP. H) Bar graphs are representing percentages of CD4^+^Rorγt^+^ Th17 in PP. Data are reported as mean±SEM. Similar results were obtained in two independent experiments with n=6. Statistical significance of each parameter was assessed by ANOVA followed by paired group comparison. *p < 0.05, **p < 0.01, ***p < 0.001, ****p < 0.0001 compared with indicated groups.

**Figure 9:**
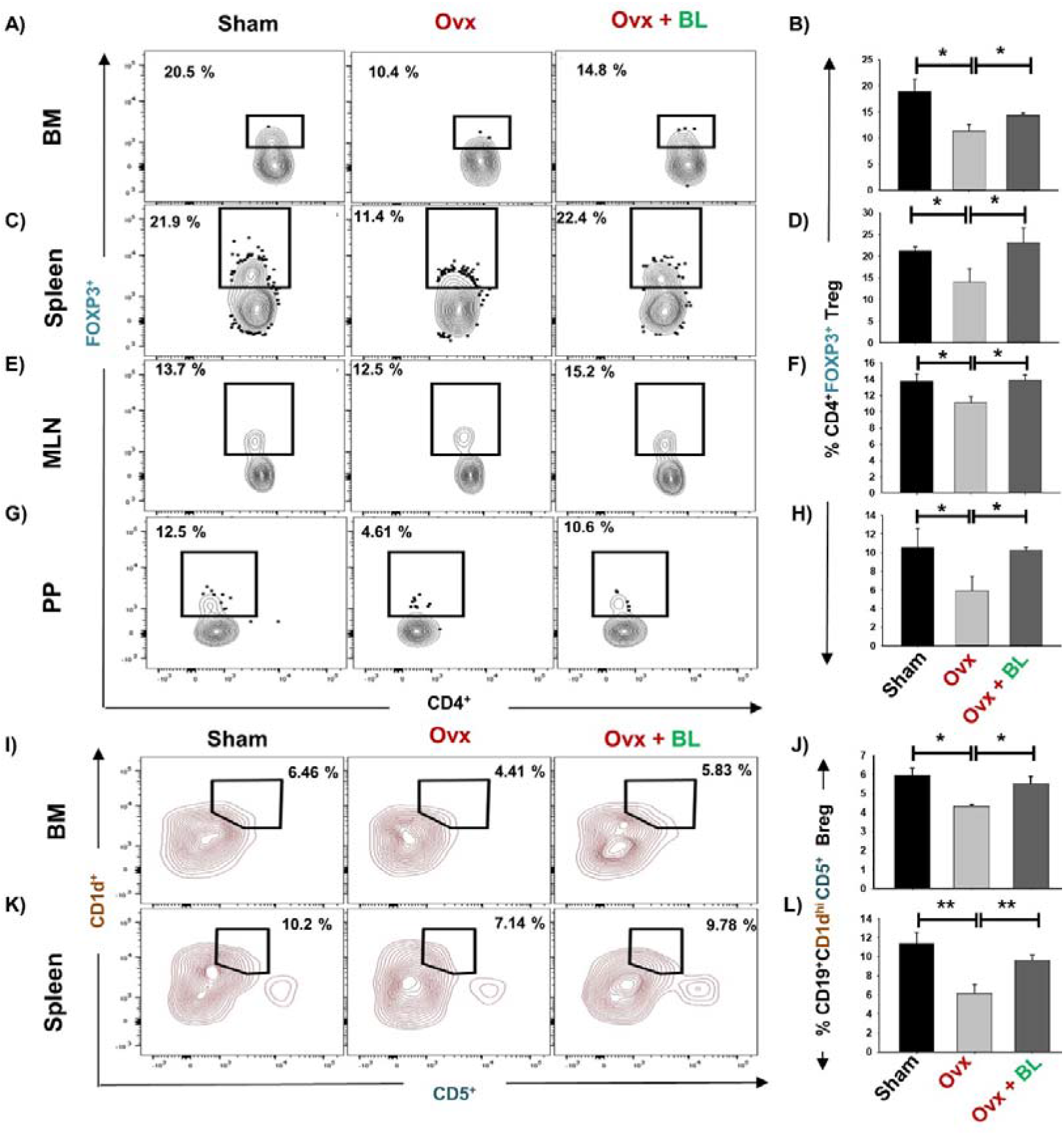
BL administration enhances anti-osteoclastogenic Tregs and Bregs *in vivo*. Cells from various lymphoid organs were harvested and analysed for CD4^+^Foxp3^+^ Tregs. A) Contour plots are representing percentages of CD4^+^Foxp3^+^ Tregs in BM. B) Bar graphs representing percentages of CD4^+^Foxp3^+^ Tregs in BM. C) Contour plots are representing percentages of CD4^+^Foxp3^+^ Tregs in spleen. D) Bar graphs representing percentages of CD4^+^Foxp3^+^ Tregs in spleen. E) Contour plots are representing percentages of CD4^+^Foxp3^+^ Tregs in MLN. F) Bar graphs representing percentages of CD4^+^Foxp3^+^ Tregs in MLN. G) Contour plots are representing percentages of CD4^+^Foxp3^+^ Tregs in PP. H) Bar graphs representing percentages of CD4^+^Foxp3^+^ Tregs in PP. I) Contour plots representing the percentages of CD19^+^CD1d^hi^ CD5^+^ Bregs in BM. J) Bar graphs representing the percentages of CD19^+^CD1d^hi^ CD5^+^ Bregs in BM. K) Contour plots representing the percentages of CD19^+^CD1d^hi^ CD5^+^ Bregs in spleen. L) Bar graphs representing the percentages of CD19^+^CD1d^hi^ CD5^+^ Bregs in spleen. Data are reported as mean±SEM. Similar results were obtained in two independent experiments with n=6. Statistical significance of each parameter was assessed by ANOVA followed by paired group comparison. *p < 0.05, **p < 0.01 and ***p < 0.001 compared with indicated groups.

### BL suppresses osteoclastogenesis in human PBMCs

Moreover, we corroborated our findings of the effect of BL in suppressing murine osteoclastogenesis in human samples. Human PBMCs when treated with BL-CM resulted in a concentration-dependent decrease in the number of multinucleated TRAP-positive cells (Fig. 11A-D). Furthermore, area measurement using an Image J software indicated significant reduction in the area of multinucleated osteoclasts in response to BL-CM (Fig. 11E). These data confirms the anti-osteoclastogenic effect of BL in human PBMCs.

**Figure 10:**
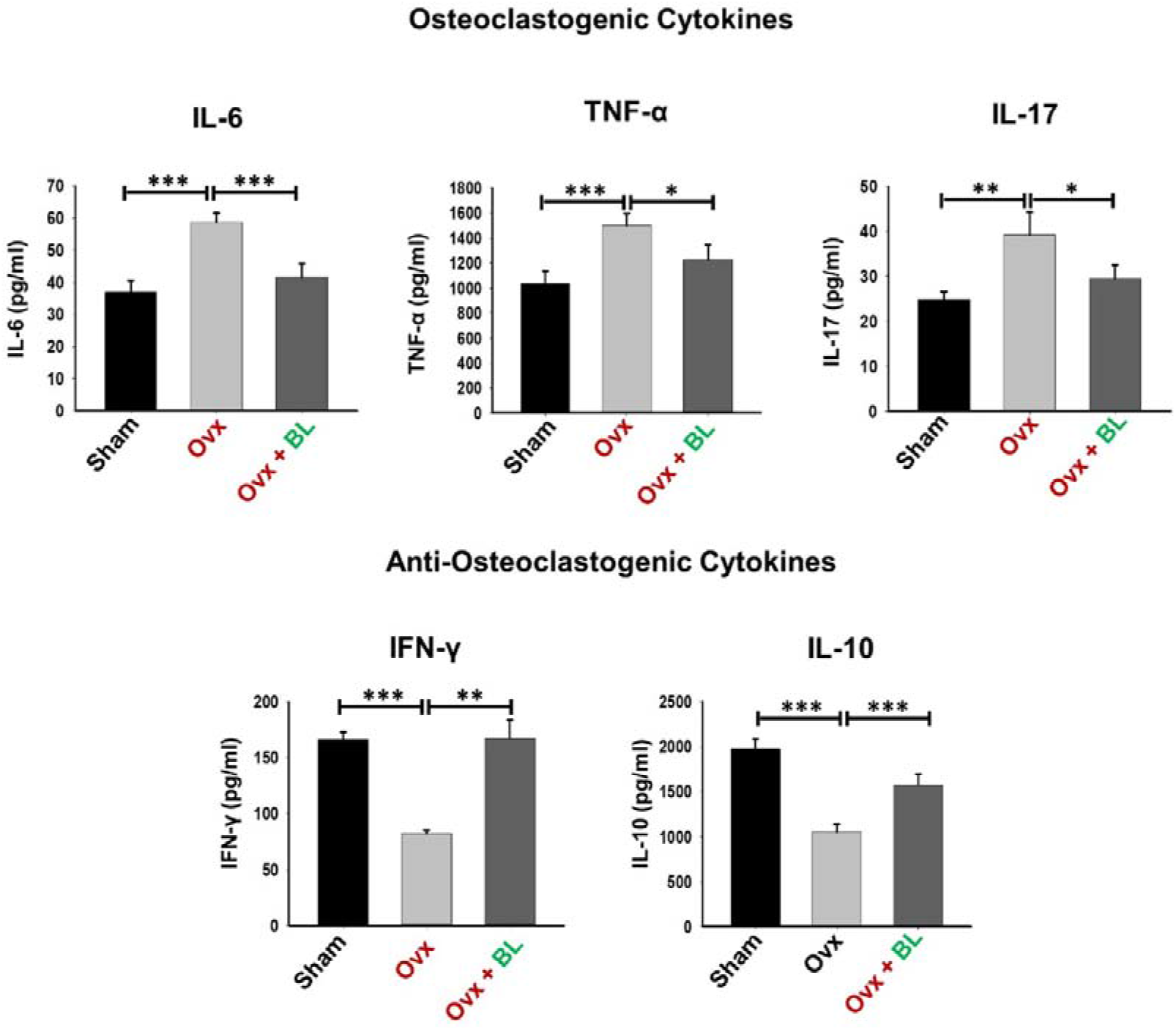
BL modulates cytokines balance in Ovx mice. Osteoclastogenic cytokines were analysed in serum samples of mice by ELISA. Anti-osteoclastogenic cytokines were analysed in serum samples of mice by ELISA. The results were evaluated by using ANOVA with subsequent comparisons by Student t-test for paired or non-paired data, as appropriate. Values are expressed as mean ± SEM (n=6) and similar results were obtained in two independent experiments. Statistical significance was defined as p≤0.05, *p≤0.05, **p < 0.01 ***p≤0.001 with respect to indicated mice group.

**Figure 11:**
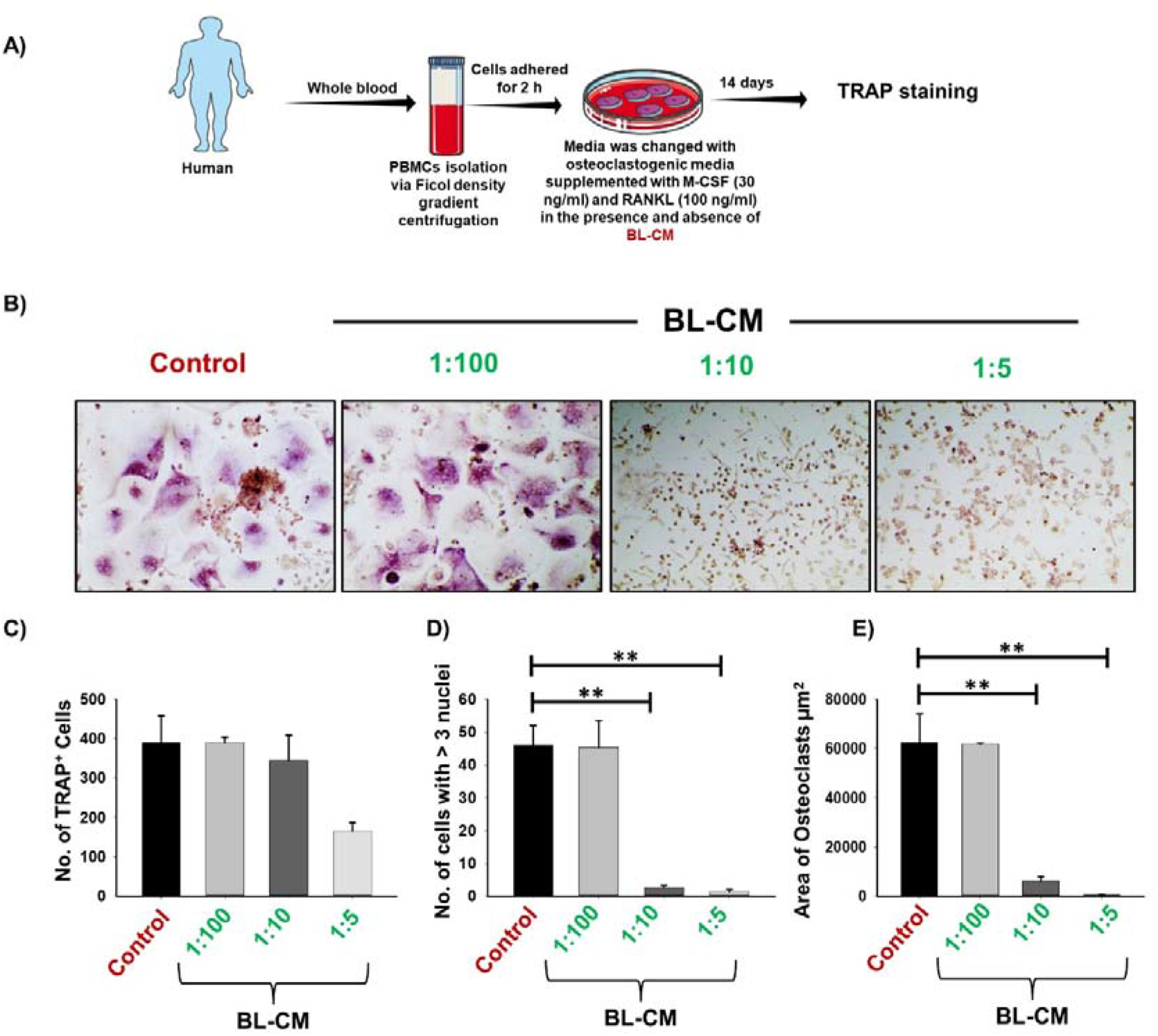
BL-CM suppress osteoclastogenesis in human PBMCs: A) Osteoclast differentiation was induced in human PBMCs with M-CSF (30 ng/ml) and RANKL (50 ng/ml) with or without *Bifidobacterium longum*— conditioned media (BL-CM) at different ratios of 1:100, 1:10 and 1:5 for 14 days. Giant multinucleated cells were stained with TRAP and cells with ≥ 3 nuclei were considered as mature osteoclasts. B) Photomicrographs at 20 × magnifications were taken. C) Number of TRAP positive cells. D) Number of TRAP positive cells with more than 3 nuclei. E) Area of osteoclasts. The above images are indicative of one independent experiment and similar results were obtained in atleast three different independent experiments. Statistical significance was considered as p ≤ 0.05 (*p ≤ 0.05, **p ≤ 0.01, ***p ≤ 0.001) with respect to indicated groups.

**Figure 12:**
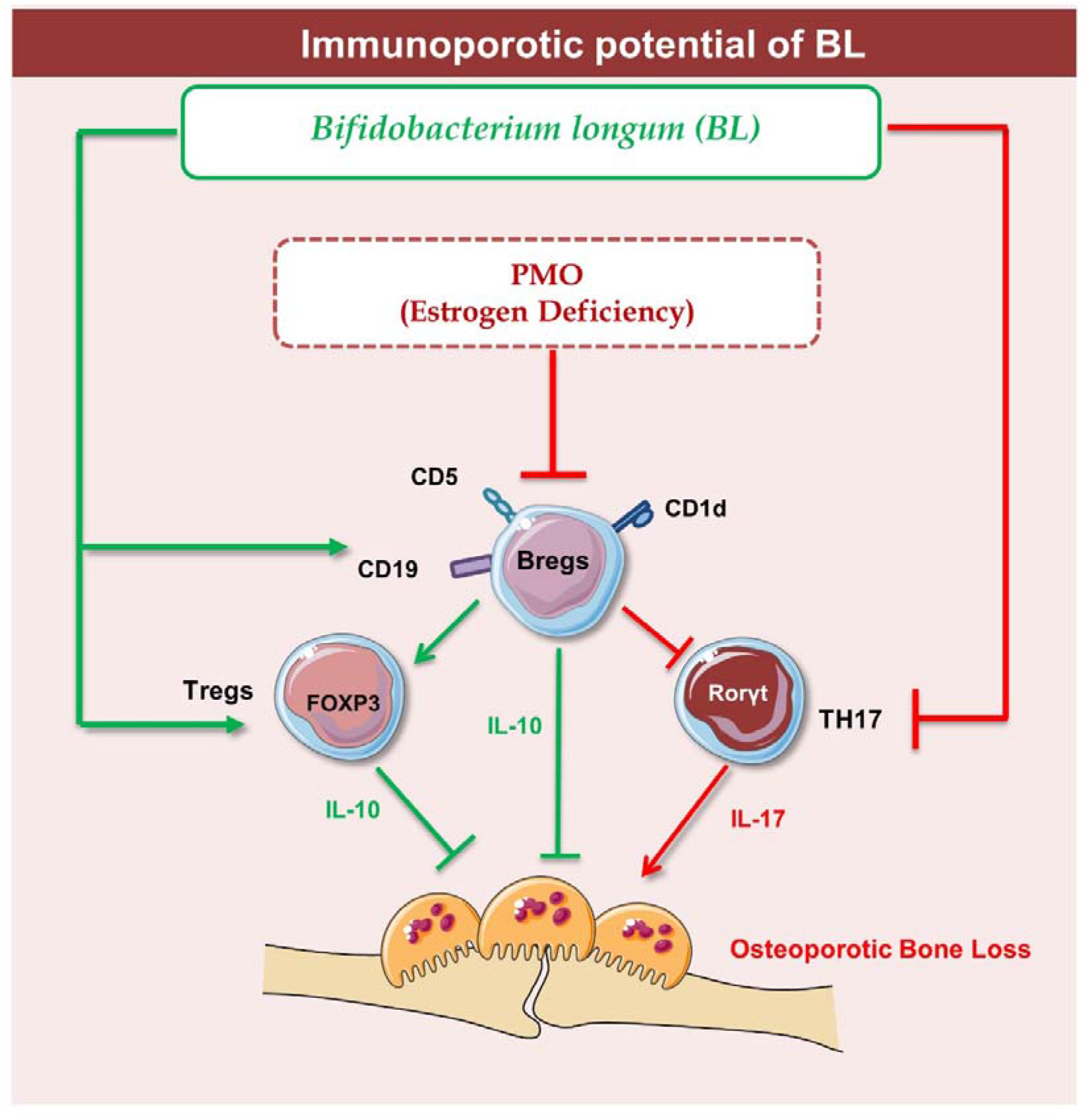
Summary of our results: Under physiological conditions, Bregs and Tregs inhibits osteoclastogenesis via producing IL-10 cytokine whereas in osteoporotic conditions, dysregulation of “Breg-Treg-Th17” cell axis promotes bone loss in osteoporotic condition. Upon BL administration, enhancement of Bregs population leads to maintenance of homeostatic balance between Tregs and Th17 cells that prevents the bone loss in osteoporotic condition (Image illustrated using Medical Art https://smart.servier.com/).

## Discussion

Osteoporosis is a chronic inflammatory condition resulting in an enhanced risk of developing fragility-related fractures at the site of the wrist, hip, and spine and is the 4^th^ most burdensome chronic disease after ischemic heart disease, dementia, and lung cancer. This skeletal disorder affects predominantly women in comparison to men and the risk increases with age. Currently, various pharmacological therapies are being employed for the treatment of osteoporosis such as bisphosphonates, denosumab, romosozumab etc. but, various health concerns have been raised with their long-term administration. A study reported that the most prescribed osteoporosis drugs such as bisphosphonates, alendronate etc. are found to be associated with the enhanced rate of developing depression and anxiety along with other adverse side effects ^18^. Thus, there is an exigent need to identify and develop safer therapies with minimal or no side effects. Experimental evidence suggests that nutritional supplementation including probiotics can maintain bone health in osteoporotic conditions. Thus, in the present study we investigated the immunoporotic potential of BL

In cultures, *Lactobacillus rhamnosus* (LR) and *Lactobacillus reuteri* suppress RANKL-mediated differentiation of osteoclasts precursors into mature osteoclasts ^19,20^. We observed that BL-CM too suppressed RANKL-induced osteoclastogenesis in murine and human precursor cells without affecting cell viability (Fig. S2). Upon adhesion to the bone surface, polarization, and re-organization of the cytoskeleton structure in osteoclasts leads to the generation of F-actin ring structure. The organization of this dynamic F-actin structure is essential for resorptive functions of osteoclasts ^21^. Our imaging data from cultured cells show that the suppression of osteoclast formation accompanied the diminished ability of osteoclasts to remain functional. These findings were corroborated in vivo as resorptive lacunae and pits measured by SEM and AFM showed significant increase in ovx mice compared with sham and BL treatment decreased it. Also, a reduction in the frequencies of osteoclast precursors in the BM (prime site of osteoclastogenesis) is further indicative of reduced osteoclasts differentiation under in vivo conditions (Fig. S3).

Under osteoporotic conditions, BL treatment resulted in bone conservation at both axial and appendicular sites and at both trabecular and cortical envelopes. Increased bone mass and cortical thickness in BL treatment appear to have contributed to increased resistance to bending failure of femur over the ovx group. An improved trabecular microarchitecture in ovx mice treated with BL over ovx is likely to confer greater resistance to compressive strength, which however, has not been measured.

Our data suggest that BL acts by direct suppression of osteoclasts as well as by immunomodulation which results in an unfavorable environment for osteoclast formation and function. Additionally, our group along with others reported that probiotics such as *Lactobacillus rhamnosus, Lactobacillus casei, Lactobacillus acidophilus, Bacillus clausii,* etc. enhance bone mass by regulating the intricate balance of “Treg-Th17” immune cells ^10,20,24^. Till date, no study has reported the immunomodulatory potential of BL in regulating bone health. Building upon this evidence, we too were interested in evaluating the immunomodulatory potential of BL. Our flow cytometric data indicate that treatment with BL-CM significantly increased the differentiation of naïve T cells into Tregs along with simultaneous inhibition of Th17 cells under in vitro conditions. In line with this, our in vivo data too confirms the immunomodulatory potential of BL in ovx mice model. Accumulating evidence suggests that under both physiological and disease conditions differentiation of Tregs and Th17 cells is regulated by Bregs via the IL-10, IL-35, and TGF-β cytokines ^25,26^. Recently, we for the first time reported that Bregs possess strong osteo-protective potential in an IL-10 dependent manner, and any decrease in the frequencies of Bregs was directly linked with pathophysiology of osteoporosis ^7^. Collectively, these studies suggest that modulation of Bregs would be a favorable therapeutic approach for the treatment of osteoporosis. Interestingly, our data from present study suggest that BL treatment significantly enhanced the differentiation of splenic B cells into IL-10^+^Bregs. Moreover, these BL stimulated Bregs significantly enhanced differentiation of Tregs along with concurrently decreasing differentiation of Th17 cells when co-cultured with BL-Bregs even under non-Treg-Th17 polarizing conditions. In addition, these BL-Bregs display enhanced anti-osteoclastogenic potential in comparison to non-BL-Bregs under in vitro conditions. Collectively, these critical experiments robustly establish the immunoporotic potential of Bregs.

Supplementation of BL in ovx mice significantly enhanced bone health via modulating the nexus between Bregs, Tregs, and Th17 immune cells. Of note, it has been observed that deficiency of estrogen hormone significantly enhanced the level of inflammatory cytokines such as IL-17 while substantially reducing the level of anti-osteoclastogenic cytokines such as IL-10 thereby augmenting bone loss in ovx mice. Importantly, our serum cytokine data further attest to the immunomodulatory potential of BL where administration of BL significantly reduced the levels of IL-17 cytokine (signature cytokine of Th17 cells) along with simultaneous induction of IL-10 cytokine (signature cytokine of Treg and Breg cells). Lastly, we decided to explore the possibility of our novel findings from pre-clinical model to clinical samples and observed that BL suppressed RANKL induced osteoclastogenesis in human PBMCs. Collectively, our present study for the first time highlights the role of BL in skeletal homeostasis and emphasizes the probiotic BL as a novel osteoprotective agent for the treatment and management of osteoporosis. BL supplementation significantly enhances BMD, strength, and micro-architecture of bones in osteoporotic mice model via modulating the “Breg-Treg-Th17” cells axis (Fig.12). However, the secretory metabolites responsible for the Immunoporotic potential of BL are still warranted, thereby opening new avenues for further research.

## Material and Methods

### Reagents and Antibodies

The following antibodies/kits were procured from eBiosciences (USA): PerCp-Cy5.5 Anti-Mouse-CD4-(RM4-5) (550954), APC Anti-Mouse/Rat-Foxp3 (FJK-16s) (17-5773), PE Anti-Human/Mouse-Rorγt (AFKJS-9) (12-6988), PerCp-Cy5.5 Anti-Mouse-CD19 (1D3) (45-0193-82), PE-Cy7 Anti-Mouse-CD5 (53-7.3) (25-0051-81), APC Anti-Mouse-CD1d (1B1) (17-0011-82), APC Cy7 Anti-Mouse-F4/80-(BM8) (47-4801-82), Foxp3/Transcription factor staining buffer (0-5523-00) and RBC lysis buffer (00-4300-54). Following ELISA kits were brought from R&D: Mouse IL-10 (M1000B) and Mouse IL-17 (M1700) Quantikine ELISA kits. The Following ELISA kits and reagents were brought from BD (USA): Mouse IL-6 (OptEIA^TM^-555240), Mouse TNF-α (OptEIA^TM^-560478). Acid phosphatase, leukocyte (TRAP) kit (387A), FITC-Phalloidin (P5282), and DAPI were purchased from Sigma (USA). Macrophage-colony stimulating factor (M-CSF) (300-25) and Receptor activator of nuclear factor κB-ligand (sRANKL) (310-01), Human TGF-β1 (AF-100-21C), Murine IL-2 (AF-212-12), Murine IL-6 (AF-216-16), Murine IL-23 (200-23), were procured from PeproTech (USA). α-Minimal essential media and RPMI-1640 were purchased from Gibco (USA). Bifido broth was procured from HiMedia Labs (Hyderabad, India). *Bifidobacterium longum* UBBL-64 was procured from Unique Biotech Ltd., Hyderabad, India.

### Animals

All in vitro and in vivo experiments were carried out in 8-10 wks old female C57BL/6 J mice. Mice were maintained under specific pathogen-free (SPF) conditions at the animal facility of All India Institute of Medical Sciences (AIIMS), New Delhi, India. Mice were fed with sterilized food and autoclaved drinking water *ad-libitum*. Mice were exposed to bilateral-ovariectomy (ovx) and sham surgery was performed on mice after anesthetizing mice with ketamine (100–150 mg/kg) and xylazine (5–16 mg/kg) intraperitoneally. Subsequently, mice were randomly allocated into three groups with 6 mice in each group i.e., sham (control); ovx and ovx + *Bifidobacterium longum* (BL).

After one-week post-surgery, ovx+BL group of mice was orally gavaged with 400 ul of BL suspension containing (10^9^ cfu) daily in drinking water for a period of 6 wks. At the end of 6 wks, animals were euthanized, and blood, bones, and lymphoid tissues were harvested for further analysis. The body weight of mice was documented at regular intervals (Day 1, Day 21, and Day 45) during the dose administration period. All the measures were performed after the due approval of the protocols submitted to the Institutional Animal Ethics Committee of AIIMS, New Delhi, India (196/IAEC-1/2019).

### *Bifidobacterium longum* (BL) bacterial culture

*Bifidobacterium longum* UBBL-64 (M1395) was cultured in Bifido broth containing 0.05 % of L-cysteine under anaerobic conditions. On the following day, overnight culture was sub-cultured into fresh Bifido broth containing freshly prepared L-cysteine (0.05%), and culture was grown until it attained log phase (OD 600nm = 0.4). Next, cells were harvested and washed with 1X PBS twice to remove the traces of Bifido broth with centrifugation at 4000 rpm for 10 min and proceeded for conditioned media preparation. CM of *B. longum* was prepared by resuspending the cells with either α-MEM or RPMI-1640 antibiotic-free media and incubated for the next 3 h at 37° C. Lastly, cell-free BL-conditioned media (CM) was collected via pelleting out the bacterial cells, pH neutralized and filtered with 0.22 µm filter. Further, BL-CM was used for all downstream assays e.g. osteoclasts and immune cells differentiation.

### Osteoclast’s differentiation and TRAP staining

For osteoclasts differentiation, mouse bone marrow cells (BMCs) were isolated from 8-12 wks old C57BL/6J mice by flushing the femoral bone with complete α-MEM media. After performing RBC lysis with 1X RBC lysis buffer, BMCs were cultured overnight in a T-25 flask in endotoxin-free complete α-MEM media (10% heat-inactivated FBS) supplemented with M-CSF at 35 ng/ml concentration. The next day, non-adherent cells were collected and seeded in 96 well plates (50,000 cells/well) in complete α-MEM media supplemented with M-CSF (30 ng/ml) and RANKL (100 ng/ml) in the presence or absence of BL-CM at different ratios viz. 1:100, 1:10 and 1:5 for 4 days. On day 3, half media was replenished with fresh complete α-MEM media supplemented with fresh factors. Lastly, for monitoring the generation of multinucleated osteoclasts, tartrate-resistant acid phosphatase (TRAP) staining was carried out according to the manufacturer’s instructions. Briefly, at the end of incubation, cells were carefully washed thrice with 1X PBS and fixed with a fixative solution containing citrate, acetone, and 3.7 % formaldehyde solution and incubated for 10 min at RT. Next, fixed cells were washed twice with 1X PBS and incubated with TRAP staining solution in dark at 37° C for 5-15 min. Multinucleated TRAP-positive cells with > 3 nuclei were considered as osteoclasts and these cells were counted and imaged using an inverted microscope (ECLIPSE, TS100, Nikon and EVOS, Thermo Scientific, USA). The area of osteoclasts was estimated with Image J software (NIH, USA).

### F-actin ring formation assay

F-actin ring formation assay was carried out as described previously ^20^. Briefly, bone marrow-derived osteoclasts precursors were seeded on glass coverslips in 12 well plates, at day 4 processing for F-actin polymerization staining was performed. Cells were washed twice with 1X PBS and fixed with 4 % paraformaldehyde (PFA) for 20 min and permeabilized with 0.1 % Triton X-100 for 5 min. Further, to block the non-specific binding, cells were blocked with 1 % BSA for 30 min and stained with FITC-labelled-phalloidin for 1 h at RT in dark. Lastly, cells were stained with DAPI (10 µg/ml) for 5 min in dark. Finally, slides were observed under an immunofluorescence microscope (Imager.Z2, Zeiss) for analyzing F-actin ring formation.

### Treg and Th17 differentiation assay

Splenic naïve T cells from C57BL/6 mice were purified by magnetic separation described previously ^7^. Briefly, after RBC lysis cells were subjected to a T cell enrichment cocktail (BD, USA) and untouched negatively selected CD4^+^CD25^-^ naïve T cells (purity > 95 %) were seeded in anti-CD3 (10 µg/ml) and anti-CD28 (2 µg/ml) mAbs coated 48 well plate. For Tregs differentiation, cells were cultured in RPMI-1640 media and stimulated with IL-2 (10 ng/ml), TGF-β1 (5 ng/ml), anti-IL-4 (5 µg/ml), and anti-IFN-γ (5 µg/ml) in the presence or absence of BL-CM at different ratios. For Th17 cell differentiation, naïve T cells were stimulated with TGF-β1 (2 ng/ml), IL-6 (30 ng/ml), IL-23 (20 ng/ml), anti-IL-4 (10 µg/ml), and anti-IFN-γ (10 µg/ml) in the presence or absence of BL-CM and incubated for 3 days. At the end of incubation, cells were harvested, and flow cytometry was performed for estimating the percentages of CD4^+^Foxp3^+^Treg and CD4^+^Roryt^+^Th17 cells.

### B cell purification and activation

Splenic B cells from C57BL/6 mice were purified by magnetic separation described previously ^7^. Briefly, following RBC lysis resulting cells were subjected to a B cell enrichment cocktail (BD, USA) and incubated for 30 min at 4°C. After washing labeled cells were incubated with streptavidin particles plus-DM for 30 min at 4° C. Further, cells underwent magnetic separation and negative fraction having B cell purity (> 95 %) were cultured in 24 well plates (2 × 10^6^) in the presence or absence of LPS (10 µg/ml) and BL-CM (1:5) for 24 h at 37°C in 5 % CO_2_ incubator. At the end of incubation, LPS and LPS + BL-CM induced Bregs were harvested and processed for either flow cytometry or co-culturing experiments (with BMCs for osteoclastogenesis and naïve T cells for Treg/Th17 cell differentiation).

### Co-culture of Bregs with BMCs and naïve T cells

For estimating the potential of BL-CM induced Bregs to suppress osteoclasts differentiation, BMCs were co-cultured with either Bregs or BL-CM induced Bregs in 96 well plates in different ratios (10:1, 5:1, and 1:1) in the presence of M-CSF (30 ng/ml) and RANKL (100 ng/ml) for 4 days. At an interval of 2 days, half media was replenished with media containing fresh factors. After 4 days of incubation, TRAP staining was performed to evaluate the generation of osteoclasts.

For evaluating the potential of Bregs to modulate Treg-Th17 differentiation, BL-CM induced Breg were co-cultured with negatively selected naïve T cells (CD4^+^CD25^-^ T cells) at 1:1 in anti-CD3 (10 µg/ml) and anti-CD28 (2 µg/ml) mAbs coated 48 well plates under non-polarization conditions. On day 4, cells were harvested, and flow cytometry was performed for estimating the percentages of CD4^+^Foxp3^+^IL-10^+^ Tregs and CD4^+^Rorγt^+^IL-17^+^ Th17 cells.

### Osteoclast differentiation from human PBMCs

PBMCs were obtained from heparinized blood by gently layering the blood on Histopaque at 1:3 in a 15 ml tube and centrifuged at 800 X g for 25 min at RT (brake off). Buffy coat layer beneath the plasma was carefully collected in a separate tube and washed with 1X PBS by inverting the tube gently and centrifuging the tube at 400 X g for 10 min. at 4°C. Obtained PBMCs were seeded in a 96-well plate at a seeding density of 1X 10^6^ cells/ well and the plate was incubated for 2 h in humidified 5 % CO2 incubator and washed with α-MEM twice (without FBS). Upon adherence, cells were incubated with α-MEM supplemented with 10 % FBS, M-CSF (30 ng/ml), and RANKL (100 ng/ml) in the presence or absence of BL-CM at different ratios (1:100, 1:10, and 1:5), and the plate was incubated for the next 14 days in a CO_2_ incubator with half media replenishment on every 3^rd^ day (i.e., 72 h). At the end of incubation, TRAP staining was performed for evaluating the differentiation of multinucleated osteoclasts. All the measures were performed after the due approval of the protocols submitted to the Institute Ethics Committee for Post Graduate Research (IECPG-482), AIIMS, New Delhi, India.

### Scanning electron microscopy (SEM)

SEM was performed for the cortical region of femoral bones as described previously ^10,20,24^. Briefly, bone samples were kept in 1%-Triton-X-100 for 2-3 days and later samples were transferred to 1X PBS buffer till the final analysis was performed. Next, after the preparation of bone slices, samples were dried under the incandescent lamp and sputter coating was done. Afterward, bones were scanned in a Leo-435-VP microscope equipped with a 35 mm photography system. SEM images were digitally photographed at 100x magnification to catch the finest cortical region. After imaging, SEM images were further analyzed by MATLAB (Math Works, Natick, MA, USA).

### Atomic force microscopy (AFM)

Upon drying femur bones with 60 W lamps for 6 h followed by high vacuum drying, samples were analyzed by Atomic Force Microscope (AFM) (INNOVA ICON, Bruker) set in Acoustic AC mode. This was assisted by cantilever (NSC 12(c) MikroMasch, Silicon Nitride Tip) and NanoDrive version 8 software, set at a constant force of 0.6 N/m with a resonant frequency at 94–136 kHz. Images were recorded at a scan speed of 1.5–2.2 lines/s in the air at room temperature. Images were later processed and analyzed by using nanoscope analysis software.

### Micro-computed tomography (µ-CT) measurements

µ-CT scanning and analysis were performed using in vivo X-ray SKYSCAN 1076 scanner (Aartselaar, Belgium) tomography. First, scanning was done at 50 kV, 204 mA using a 0.5 mm aluminium filter by positioning samples at the right orientation in the sample holder. For the reconstruction process, NRecon software was employed. After reconstruction, ROI was drawn at a total of 100 slices in secondary spongiosa at 1.5 mm from the distal border of growth plates and further processed for CTAn software for evaluating and calculating micro-architectural parameters of bone samples. Several 3D-histomorphometric parameters were obtained viz. Bone volume/Tissue volume (BV/TV), Trabecular thickness (Tb.Th), Trabecular separation (Tb.Sp) etc. The volume of interest of u-CT scans made for trabecular and cortical regions was used to determine the BMD of LV5, femur, and tibia. BMD was measured by using hydroxyapatite phantom rods of 4 mm diameter with known BMD (0.25 g/cm3 and 0.75 g/cm^3^) as a calibrator (Sapra 2021).

### Bone strength testing

To measure the bio-mechanical properties of bones, femoral bone of mice in all the respective groups were exposed to three-point bending by employing bone strength tester model TK-252C/RDT (Muromachi Kikai Co. Ltd., Tokyo, Japan). For this, femoral bone was placed on the two supports that was kept at a constant distance of 1 cm. By means of load displacement curves, following bone mechanical strength parameters was evaluated : maximum power (N), energy to fracture (mJ) and stiffness (N/mm).

### Flow cytometry

Cells were harvested from various lymphoid organs (viz. BM, Lymph nodes, Spleen & PPs) and stained with antibodies specific for macrophages, Bregs, Tregs, and Th17 cells. For the macrophage panel, BM cells were stained with anti-F4/80-APC Cy7 antibody. For Breg cell panel, cells were stained with anti-CD19-PerCP-Cy5.5, anti-CD5-PE-Cy7, and anti-CD1d-APC antibodies. For Treg/Th17 cells were first stained with anti-CD4-PerCP-Cy5.5 antibody for cell surface staining and incubated for 30 min in dark on ice. After washing, cells were further fixed and permeabilized with a 1X fixation-permeabilization buffer for 30 min on ice in dark. Finally, for intracellular staining, cells were stained with anti-Rorγt-PE, anti-Foxp3-APC, anti-IL-10-BV421 and anti-IL-17-APC for 45 min. After washing, cells were acquired on BD-LSR Fortessa (USA). Flowjo-10 (TreeStar, USA) software was used to analyze the samples, and the gating strategy was done as per experimental requirements.

### Enzyme-linked immunosorbent assay (ELISA)

ELISA was carried out for the quantitative assessment of following cytokines i.e., IL-6, IL-10, IL-17, TNF-α, and IFN-γ in blood sera of all the mice groups by utilizing commercially available kits as per the manufacturer’s instructions.

### Statistical analysis

Statistical differences between the distinct groups were evaluated by employing analysis of variance (ANOVA) with subsequent analysis via student t-test paired or unpaired as appropriate. All the values in the data are expressed as Mean ± SEM (n=6). Statistical significance was determined as p≤0.05 (*p < 0.05, ** p < 0.01, *** p, 0.001) with respect to the indicated groups.

## Acknowledgment

This work was financially supported by the following projects: DST-SERB (EMR/2016/007158), Govt. of India and intramural project from All India Institute of Medical Sciences (AIIMS, AI-798), New Delhi-India sanctioned to RKS. LS, NS, AB, PKM and RKS acknowledge the Department of Biotechnology AIIMS, New Delhi-India for providing infrastructural facilities. LS thank UGC, NS thank DBT, and AB thank DST SERB for research fellowship.

## Author contributions

RKS contributed in conceptualization and investigation of study. LS, NS, CS and AB contributed to methodology and formal analysis of data; KG helped in µ-CT analysis; MM performed osteoclast culture in human PBMCs; LS developed ovx mice model; PKM carried out cytokine analysis; NC contributed in µ-CT and bone mechanical strength data analysis. RKS and LS contributed in writing and editing of manuscript. BV provided the valuable inputs. All authors reviewed the manuscript.

## Conflicts of Interest

The authors declare no conflicts of interest.

## Compliance with ethical standards

All applicable institutional and/or national guidelines for the care and use of animals and human samples were followed.

---

**Fig S1:**
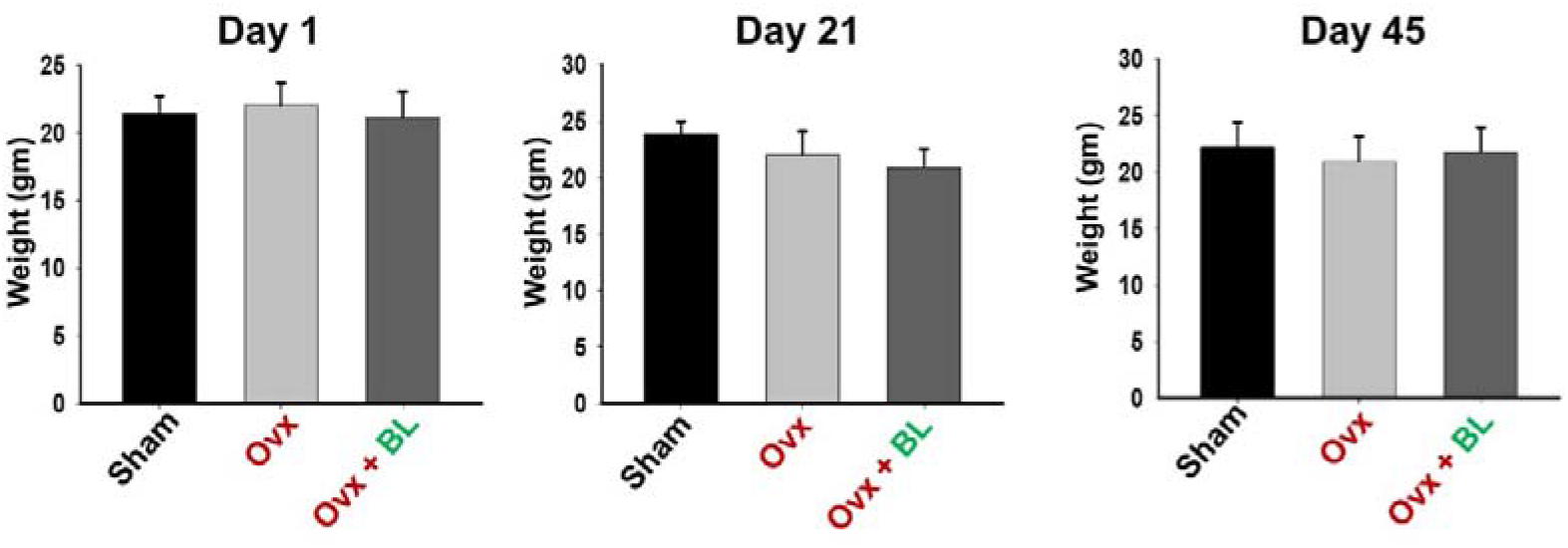
Body weight of Mice. Mice were divided into 3 groups viz. Sham, Ovx and Ovx + BL group that received BL at 10^9^ CFU/day orally reconstituted in drinking water. At the end of 45 days, mice were sacrificed and analysed for various parameters and body weight was monitored at regular intervals; Values are reported as mean ± SEM (n=6).

**Fig S2:**
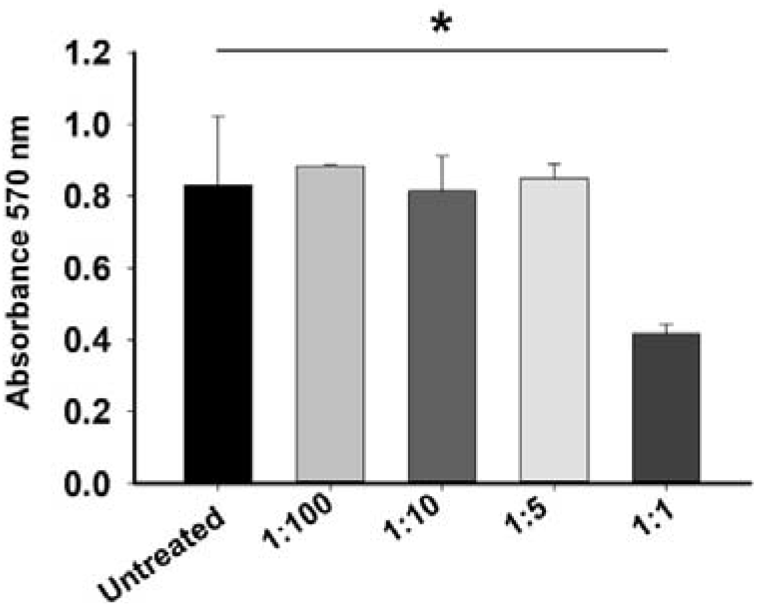
Cell cytotoxicity assay. Bone marrow cells (BMCs) were treated with different dilutions of BL supernatant for 48 h and MTT assay was performed for evaluating the cell cytotoxic effects of BL.

**Fig S3:**
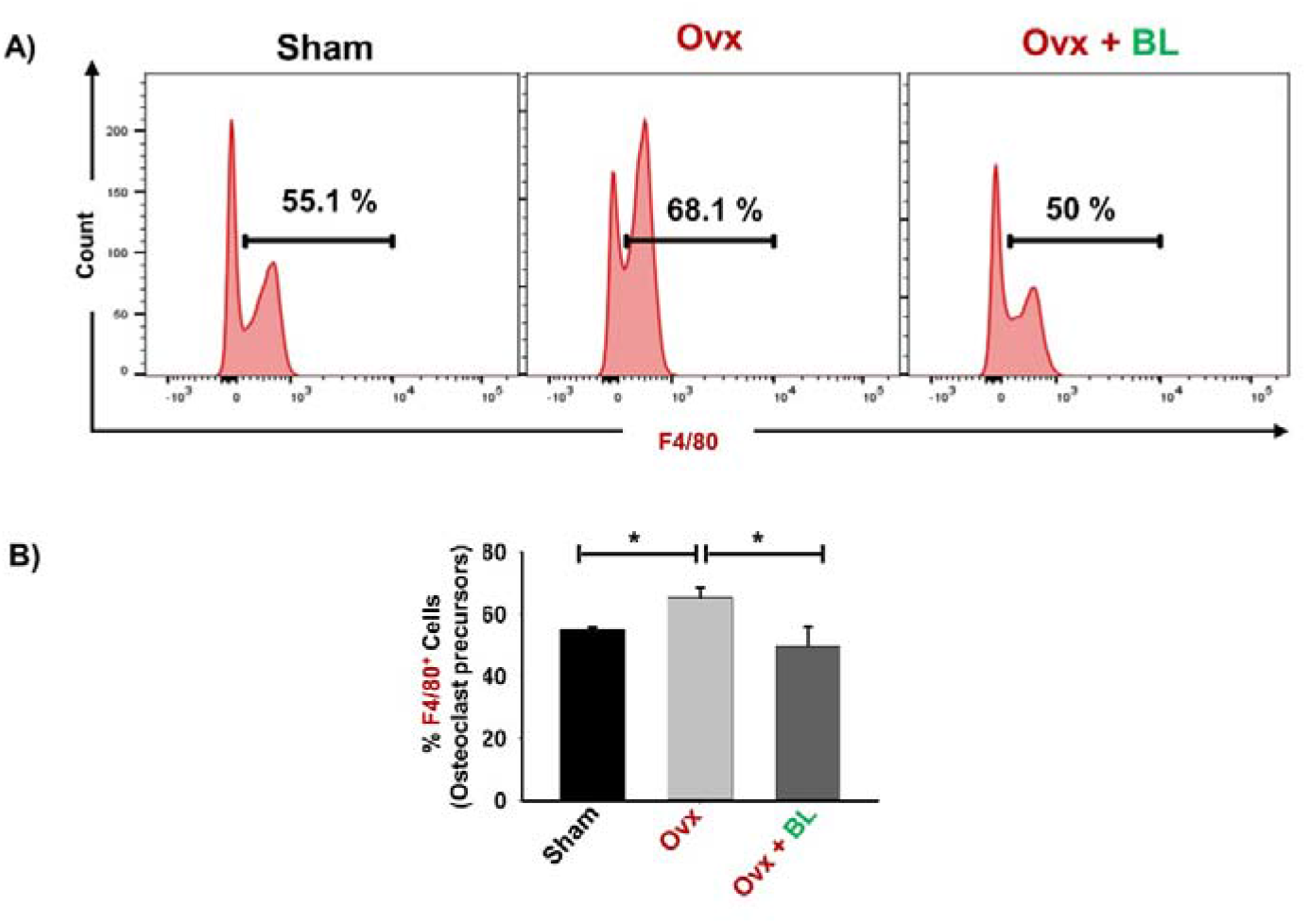
BL administration reduces F4/80^+^ osteoclasts precursors in BM. A) Histograms representing percentages of macrophages in all the groups B) Graphical representation of percentages of macrophages in three groups. The results were evaluated by using ANOVA with subsequent comparisons by Student t-test for paired or non-paired data, as appropriate. Values are expressed as mean ± SEM (n=6) and similar results were obtained in two independent experiments. Statistical significance was defined as p≤0.05, *p≤0.05, **p < 0.01 ***p≤0.001 with respect to indicated mice group.

## Notes

### Competing Interest Statement

The authors have declared no competing interest.

